# HIV-1 co-receptor usage and variable loop contact impacts V3 loop bnAb susceptibility

**DOI:** 10.1101/568469

**Authors:** Ludy Registre, Yvetane Moreau, Sila Toksoz Ataca, Surya Pulukuri, Timothy J. Henrich, Nina Lin, Manish Sagar

## Abstract

In clinical trials, HIV-1 broadly neutralizing antibodies (bnAbs) effectively lower plasma viremia and delay virus reemergence after antiretroviral treatment is stopped among infected individuals that have undetectable virus levels. Presence of less neutralization susceptible strains prior to treatment, however, decreases the efficacy of these antibody-based treatments. The HIV-1 envelope glycoprotein harbors extensive genetic variation, and thus, neutralization sensitivity often cannot be predicted by sequence analysis alone. Sequence-based prediction methods are needed because phenotypic-based assays are labor intensive and not sensitive. Based on the finding that phenotypically confirmed CXCR4- as compared to exclusive CCR5-utilizing strains are less neutralization sensitive, especially to variable loop 1 and 2 (V1-V2) and V3 loop bnAbs, we show that an algorithm that predicts receptor usage identifies envelopes with decreased V3 loop bnAb susceptibility. Homology modeling suggests that the primary V3 loop bnAb epitope is equally accessible among CCR5- and CXCR4-using strains although variants that exclusively use CXCR4 have V3 loop protrusions that interfere with CCR5 receptor interactions. On the other hand, homology modeling also shows that envelope V1 loop orientation interferes with V3 loop directed bnAb binding, and this accounts for decreased neutralization sensitivity in some but not all cases. Thus, there are likely different structural reasons for the co-receptor usage restriction and the differential bnAb susceptibility. Algorithms that use sequence data to predict receptor usage and antibody-envelope homology models can be used to identify variants with decreased sensitivity to V3 loop and potentially other bnAbs.

**AUTHOR SUMMARY:** HIV-1 broadly neutralizing antibody (bnAb) therapies are effective, but the pre-existence of less susceptible variants may lead to therapeutic failure. Sequence-based methods are needed to predict pre-treatment variants’ neutralization sensitivity. HIV-1 strains that use the CXCR4 as compared to the CCR5 receptor are less neutralization susceptible, especially to V1-V2 and V3 loop bnAbs. A sequence-based algorithm that predicts receptor usage can identify envelope variants with decreased V3 loop bnAb susceptibility. While the inability to utilize the CCR5 receptor maps to a predicted protrusion in the envelope V3 loop, this viral determinant does not directly influence V3 loop bnAb sensitivity. Furthermore, homology modeling predicted contact between the envelope V1 loop and an antibody also impact V3 loop bnAb susceptibility in some but not all cases. An algorithm that predicts receptor usage and homology modeling can be used to predict sensitivity to bnAbs that target the V3 loop and potentially other envelope domains. These sequence-based methods will be useful as HIV-1 bnAbs enter the clinical arena.

## INTRODUCTION

Multiple broadly neutralizing antibodies (bnAbs) are being examined as novel therapeutics against human immunodeficiency virus type 1 (HIV-1) infection [1–6]. In contrast to the current highly effective antiretroviral drugs (ARVs), antibody-based therapies require less frequent dosing, can be effective against drug resistant variants, and may potentiate host humoral responses [7]. Prior to initiating ARVs, HIV-1 infected patients are routinely evaluated for the presence of drug resistant strains, primarily using sequence-based methods [8]. Sequence-based methods are also needed to identify pre-treatment variants with reduced bnAb susceptibility because phenotypic-based assays are cumbersome and lack sensitivity [1,4].

BnAbs attach to diverse envelope (Env) domains, such as the apex, high mannose patch, CD4 binding site (bs), surface unit (gp120) – transmembrane (gp41) interface, and gp41 membrane proximal external region (MPER) [9]. The apex is targeted by variable loop 1 and 2 (V1-V2) directed bnAbs that bind the asparagine (N)-linked glycan at Env position 160 (N160) [10]. Anti-variable loop 3 (V3 loop) bnAbs attach to an N-linked glycan at Env position 332 in the high mannose patch [11]. While the activity of these bnAbs primarily depends on the presence of these glycans, other amino acids, especially those in and around the V1-V2 and V3 Env regions, also impact neutralization [12,13]. In addition to being antibody targets, the V1-V2 and V3 loops also influence binding to either the CCR5 or CXCR4 co-receptor, and this attachment is necessary for host cell entry [14–16]. This overlap provides the scientific basis for speculating that there is an association between the receptor a virus utilizes to enter cells and its bnAb sensitivity. We and others have previously shown that HIV-1 subtype C (HIV-1C) and HIV-1 subtype D (HIV-1D) variants that exclusively utilize the CXCR4 receptor (termed X4) often have insertions or basic amino acid substitutions in the V3 loop compared to the strains that either only use CCR5 (classified as R5) or those that can employ either receptor (termed R5X4) [14,15]. Our group and others have also argued that certain antibody responses may select for CXCR4-utilizing variants [17–19]. Even though, the sequence signatures associated with X4 strains do not involve the primary V1-V2 and V3 bnAb epitopes, associated sequence modifications and possible linked structural changes may impact bnAb susceptibility. Both V3 loop sequence motifs and structure-based models have been used to predict CCR5 and CXCR4 usage, and thus, similar methods could potentially be used to speculate about bnAb sensitivity [20–22]. In this study, we show that co-receptor usage prediction can be used to identify Envs with decreased susceptibility to V3 loop bnAbs. Furthermore envelope-antibody homology models predict antibody sensitivity in some cases, and these types of sequence-based methods could be used to predict bnAb susceptibility in the future.

## Results

### CXCR4- as compared to CCR5-using strains are more neutralization resistant

Observing an association between co-receptor usage and bnAb susceptibility could be useful in developing sequence-based methods to determine pre-treatment bnAb sensitivity because there are numerous algorithms that predict receptor utilization based on input sequence alone. Some prior studies have suggested that HIV-1B CXCR4- as compared to CCR5-using viruses are more neutralization susceptible [23–27]. If true, CXCR4-utilizing strains should predominantly exist in individuals that have decreased neutralization capacity. We tested this prediction by comparing neutralization breadth and potency among plasma samples collected from individuals with previously well-characterized virus population [28]. Neutralization responses were compared among eleven individuals with no evidence of CXCR4-using virus (classified as R5 only) and seven subjects with a mixture of CXCR4 and CCR5 utilizing variants (termed dual-mixed (DM)) (Table S1). Samples were classified either as R5 only or as DM if the absence or the presence of CXCR4-using virus was confirmed by a bulk Env phenotype assay and in some cases by the analysis of individual Envs isolated using single genome amplification (SGA) also [28]. Samples’ neutralization capacity was assessed against eleven R5 Envs of varying subtypes because responses against this global reference panel have been previously used to estimate neutralizing breadth and potency [29,30]. Heat maps depicting the neutralization responses against the eleven Envs revealed that the individuals from the two groups were not qualitatively different (Fig. 1A). Plasma neutralization capacity was estimated using a previously defined breadth and potency (BP) score [30]. Briefly, BP score consisted of the average log normalized percent neutralization at the highest tested plasma dilution (1:50) across all the viruses in the panel. DM plasma had similar BP as compared R5 only plasma (p = 0.37) (Fig 1B). The percentage of the eleven Env reference panel neutralized at greater than 50% (termed breadth) was also similar between the DM as compared to the R5 only samples (p = 0.88) (Fig 1C). Interestingly, DM plasma (4102 and 1239) with demonstrated X4 variants (Table S1) had some of the most potent (BP score 0.92 and 0.57) and greatest neutralization breadth (100% and 82%) (Fig. 1A).

**Fig. 1.**
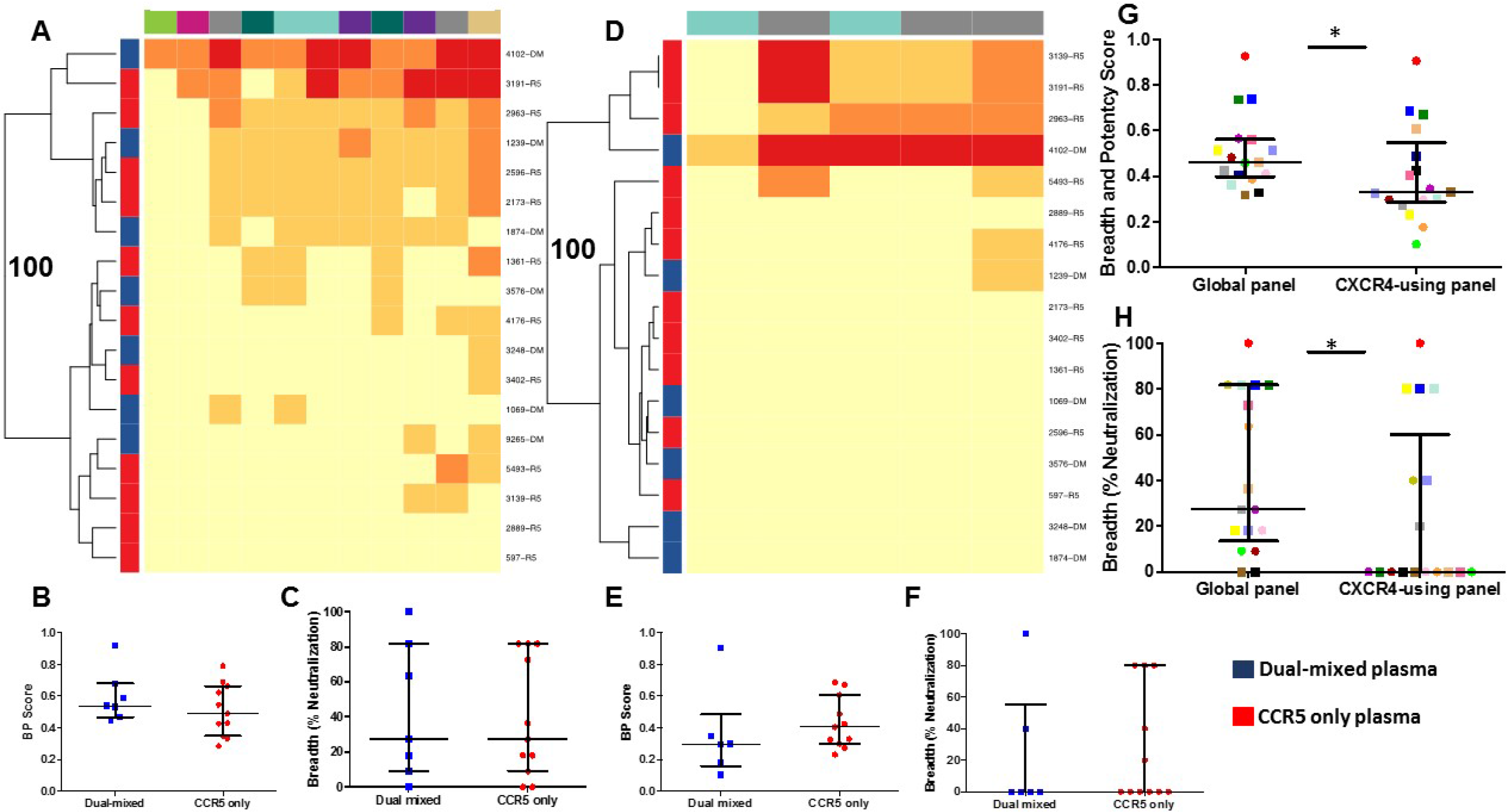
Samples containing CXCR4-using or those with CCR5 only viruses have similar neutralization potency and breadth but plasmas have decreased ability to neutralize CXCR4-using as compared to R5 strains. **(A and D)** Heatmaps show plasma neutralization against the R5 global reference Env panel **(A)** and CXCR4-using Env collection **(D)**. Each square in the heat map represents the average percent neutralization for each Env-plasma combination tested: <50% (yellow); 50-70% (light orange); 70-90% (dark orange); >90% (red). On the left, blue and red denotes DM and R5 only plasma respectively, and individual plasma IDs are listed on the right. Env subtypes above the heatmaps are indicated by color: A (khaki); B (gray); C (teal); G (green); AC (pink); CRF01_AE (dark green); and CRF07_BC (purple). The branches show the hierarchical clustering with bootstrap probability for 100 iterations. **(B and E)** Breadth and potency (BP) score for DM (blue) and CCR5 only plasma (red) against the global Env panel **(B)** and CXCR-using Env collection **(E)**. **(C and F)** Breadth (% of Envs neutralized at greater than 50% at the highest tested plasma dilution) observed for the DM (blue) and CCR5 only plasma (red) against the global Env panel **(C**) and CXCR4-using Env collection **(F)**. Comparisons done using Wilcoxon rank -sum test. **(G and H)** Plasma BP score **(G)** and breadth **(H)** against the global as compared to the CXCR4-using Env panel. In G and H, each unique plasma sample is denoted by a different color/symbol. Comparisons done using matched-pairs Wilcoxon rank-sum test. In all box plots, values are a mean from a minimum of 2 independent assays. In each box plot, lines denotes median and interquartile range. Stars denote p-value less than 0.05.

It is possible that we may have failed to observe a difference among the DM as compared to the R5 samples because of the relatively small sample size. In our analysis there was 80% power, at type 1 error level of 0.05, to detect around 1.5 fold or greater difference in the median BP score based on the observed distributions. DM as compared to the R5 only plasma may have also potentially failed to demonstrate neutralization capacity differences because the global reference panel contained R5 Envs only [29]. Neutralization capacity was examined against another Env collection consisting of CXCR4-using variants (Table S2) [14,31]. DM and R5-only plasma had similar neutralization fingerprints against this CXCR4-using Env collection (Fig 1D), and there was no significant difference in BP score (p = 0.22) (Fig 1E) or breadth (p = 0.83) (Fig 1F). In aggregate, HIV-1B CXCR4-using viruses do not predominantly exist among individuals with weak humoral immune responses.

Studies from our group and others have argued that CXCR4-using variants may emerge as a consequence of antibody pressure because X4 as compared to R5 Envs are more neutralization resistant [17–19]. We compared the eleven R5 only and seven DM plasma’s neutralization capacity against Envs in the reference global panel (all R5) to those in the CXCR4-using collection to provide further generalizability for this conclusion. The eighteen plasma had decreased neutralization capacity (around 1.5 fold lower BP score) against the CXCR4-using as compared to against the global reference Env panel (p = 0.01) (Fig 1G). Furthermore, neutralization breadth was significantly lower against the CXCR4-using as compared to the global reference Env collection (p = 0.02) (Fig 1H). These observations suggest that, in general, HIV-1B CXCR4- as compared to CCR5-using viruses are more neutralization resistant. Thus, there is an association between co-receptor usage and neutralization sensitivity.

### X4 variants are less neutralization susceptible to V1-V2 and V3 bnAbs

The linkage between co-receptor usage and neutralization susceptibility was examined in further detail by comparing available bnAb IC_50_s among variants with phenotypically-confirmed co-receptor usage in the Los Alamos CATNAP database [32]. A larger proportion of the CXCR4-using as compared to R5 variants had an IC_50_ above the highest tested bnAb concentration for the V3 directed glycan bnAb, PGT121 (p < 0.0001), the V1-V2 glycan dependent bnAbs, PG9 (p = 0.02) and PG16 (p = 0.03), but not for the CD4bs bnAb, VRC01 (p = 0.95). The tested but undetectable IC_50_s were given a value of 100 for the subsequent statistical comparisons. The CXCR4-utilizing variants had significantly higher IC_50_ to PGT121, as compared to the R5 strains (p = 0.0002) (Fig. 2A). The R5 variants also had significantly lower IC_50_s to PG9 (p = 0.03) and PG16 (p = 0.04) as compared to the CXCR4-using strains (Fig. 2B and 2C). On the other hand, CXCR4-using variants and R5 had similar neutralization susceptibility to VRC01 (p = 0.26) (Fig. 2D). Meaningful comparisons could not be done for another V3 (10-1074) and a MPER (10E8) bnAb because of low number of CXCR4-using variants with available IC_50_ data in the CATNAP database (n = 9 each). These observed neutralization differences to PGT121, PG9, and PG16 remained statistically significant even if R5 variants were compared to X4 strains only. A small number of R5X4 variants with available IC_50_ data precluded their examination as an independent group.

**Fig. 2.**
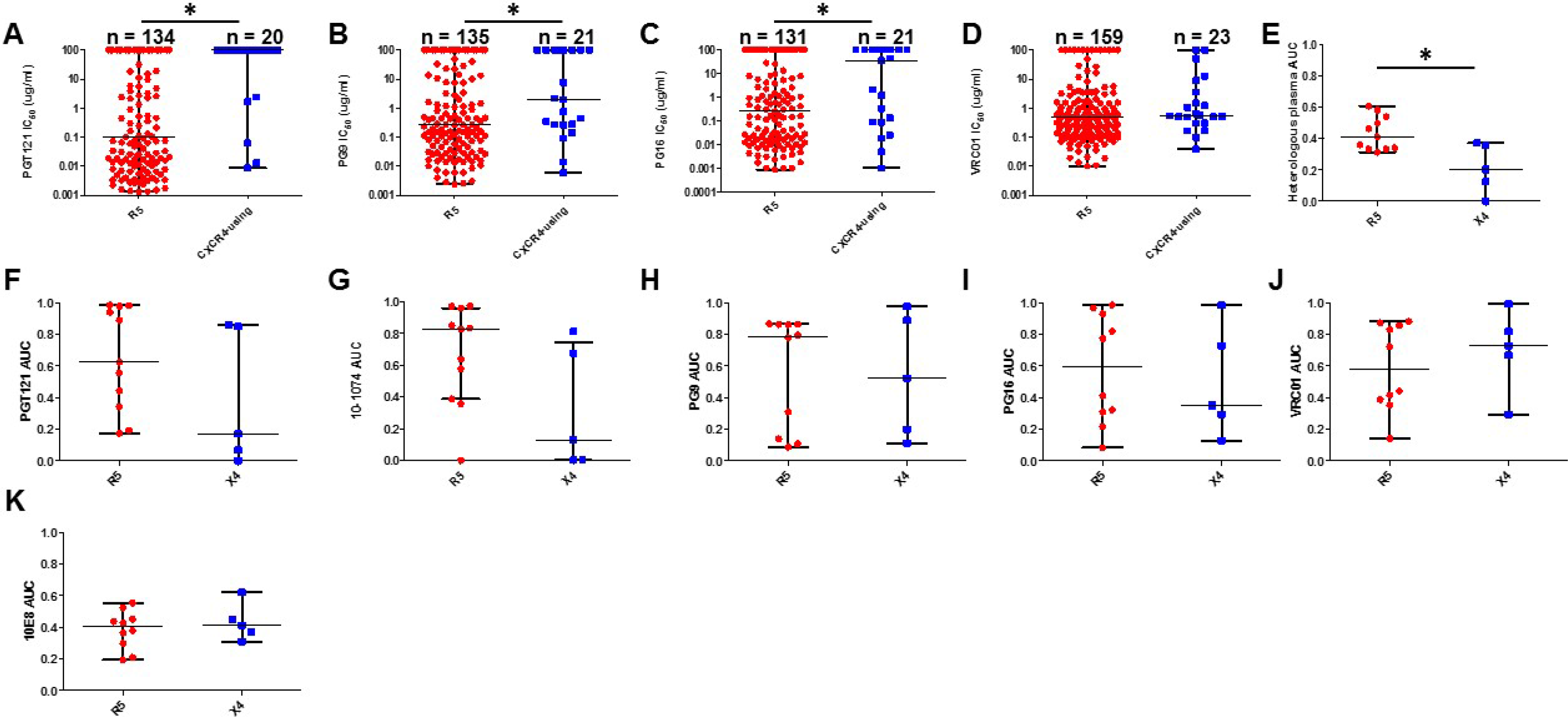
CXCR4-using as compared to R5 strains are less susceptible to heterologous antibodies and V1-V2 and V3 directed bnAbs. **(A – D)** Neutralization IC_50_s available in the Los Alamos CATNAP database among R5 (red) and CXCR4-using (blue) strains against PGT121 **(A)** PG9 **(B)**, PG16 **(C)**, and VRC01 **(D)**. In these analyses, a variant with an IC_50_ above the highest tested bnAb dilution was assigned a value of 100. **(E)** Neutralization area under the curve (AUC) (y-axis) among primary R5 (red) and X4 (blue) Env against a heterologous plasma pool. **(F – K)** Neutralization AUC (y-axis) for primary R5 (red) and X4 (blue) Envs against PGT121 **(F)**, 10-1074 **(G),** PG9 **(H)**, PG16 **(I)**, VRC01 **(J)**, and 10E8 **(K)**. Each point denotes a unique Env and the value represents mean from duplicate independent experiments. Among all the dot plots **(A – J)**, lines denotes median and interquartile range. All comparisons done using Wilcoxon rank-sum test. Stars denote p-value less than 0.05.

The CXCR4-using variants available in the Los Alamos database are often lab-adapted strains, and neutralization characteristics can change with lab passaging [33,34]. Thus, neutralization susceptibility was compared among primary Envs with phenotypically determined co-receptor usage. A total of 929 individual Envs (median = 16 Envs per subject, range= 1-239) were isolated using SGA from 33 previously classified DM anti-retroviral naïve patient samples [28,35]. A web-based prediction tool, either WebPSSM or geno2pheno, was used to predict the co-receptor usage of the isolated Envs based on the V3 loop sequence [21,22]. Some of the SGA Envs from 22 individuals were genotypically predicted to use CXCR4, and predicted CXCR4-utilizing Envs were likely not isolated from the other eleven previously characterized DM samples because of relatively limited SGA sampling compared to the bulk PCR analysis. Receptor usage phenotype was examined for some viruses that incorporated SGA isolated Envs from the seventeen samples containing sequence predicted CXCR4-utilizing strains (Table 1). Five (4102, 1239, 3248, 1924, and 3576) contained X4 Envs, and a primary X4 Env group was generated by randomly selecting one variant from each of the 5 individuals (Table S3). The Envs examined from the remaining twelve samples were either R5, even though some of them were predicted by sequence to use the CXCR4 receptor, or a mixture of R5X4 and R5 (Table 1). A primary R5 Env group was also generated by randomly selecting 1 phenotypically confirmed CCR5 only using strain from 11 different individuals (Table S3).

**Table 1.** Subject envelope characteristics.

| Subject | # of SGA Envs isolated | # of SGA Envs with predicted genotype |  | # of Envs phenotyped | # of recombinant viruses with confirmed phenotype |  |  |
| --- | --- | --- | --- | --- | --- | --- | --- |
|  |  | R5 | CXCR4 |  | R5 | X4 | R5/X4 |
| 4102 | 61 | 52 | 9 | 19 | 13 | 5 | 1 |
| 1239 | 239 | 234 | 5 | 12 | 9 | 1 | 2 |
| 3248 | 16 | 0 | 16 | 9 | 0 | 9 | 0 |
| 1924 | 27 | 24 | 3 | 2 | 1 | 1 | 0 |
| 3576 | 30 | 0 | 30 | 6 | 0 | 6 | 0 |
| 3131 | 17 | 6 | 11 | 7 | 7 | 0 | 0 |
| 1069 | 10 | 0 | 10 | 9 | 9 | 0 | 0 |
| 2327 | 6 | 0 | 6 | 4 | 1 | 0 | 3 |
| 0229 | 40 | 5 | 35 | 7 | 3 | 0 | 4 |
| 1045 | 15 | 5 | 10 | 5 | 5 | 0 | 0 |
| SC | 11 | 7 | 4 | 6 | 5 | 0 | 1 |
| 1389 | 14 | 0 | 14 | 7 | 1 | 0 | 5 |
| 1486 | 15 | 7 | 8 | 5 | 5 | 0 | 0 |
| 3026 | 110 | 109 | 1 | 4 | 4 | 0 | 0 |
| 1233 | 98 | 72 | 26 | 10 | 7 | 0 | 0 |
| 9265 | 14 | 0 | 14 | 14 | 0 | 0 | 14 |
| 1874 | 11 | 2 | 9 | 5 | 5 | 0 | 0 |

Neutralization susceptibility was compared between the group of primary X4 and R5 Envs to a standard comparator generated by pooling plasma from ten HIV-1B infected individuals, different from the seventeen subjects above. We chose to compare the groups using neutralization area under the curve (AUC) rather than the concentration required to achieve 50% inhibition (IC_50_) because the two estimates are highly correlated and AUC can be used when 50% inhibition is not observed at the highest tested concentration [30,36] (Fig. S1). The HIV-1B primary X4 Envs had a two-fold lower AUC as compared to the R5 group (p = 0.03) (Fig 2E). The primary X4 as compared to R5 variants were around 4 to 8 fold less sensitive to V3 directed antibodies PGT121 and 10-1074 although the differences only showed a statistical trend likely due to small sample size (p = 0.09 for both) (Fig 2F and 2G). The primary X4 viruses were around 2 fold less sensitive to V1-V2 antibodies, PG9 and PG16, but these differences were also not statistically significant (Fig. 2H and 2I). In contrast, the R5 as compared to the X4 variants had comparable sensitivity to CD4bs and MPER bnAbs (Fig 2J and 2K). Thus, similar to our HIV-1C findings and Los Alamos CATNAP database analysis, primary HIV-1B X4 as compared to R5 variants have decreased neutralization susceptibility, especially to V3 and possibly V1-V2 directed antibodies [18]. Thus, algorithms that predict co-receptor usage can potentially be utilized to estimate V3 and potentially V1-V2 bnAb sensitivity.

### V3 loop protrusions impact CCR5 receptor interactions but not access to the V3 loop bnAb epitope

Modification of the glycosylated amino acid at Env position 332 is the primary resistance determinant for V3 directed bnAb, but decreased sensitivity can arise even if this epitope remains intact [3,12]. All the R5 and X4 variants that were compared for their neutralization susceptibility to PGT121 and 10-1074 (Fig. 2F and 2G) had a predicted glycan at position 332. A sequence alignment of 22 X4, 31 R5X4, and 77 R5 primary Envs with phenotypically confirmed receptor usage revealed that all X4 variants, except 1924, contained a 2 to 3 amino acid V3 loop insertion either directly before the glycine (G) – proline (P) - G crown or toward the base of the V3 loop (Fig. S2). These specific V3 modifications were not found in any of the phenotypically confirmed R5 Envs. In contrast, the X4 variant in subject 1924 contained a positively charged amino acid substitution at V3 loop position 25. This change has previously been associated with CXCR4 receptor usage, although this sequence motif is not as highly predictive for exclusive CXCR4 usage as the V3 loop insertion [14,21].

We hypothesized that this observed V3 loop sequence motif associated with X4 strains likely restricted both co-receptor usage and access the primary V3 loop bnAb epitope, namely the glycan at position 332. Structural homology models were used to assess this premise. The predicted V3 loop structure of X4 Envs (1239, 1924, 4102, 3248 and 3576) was either compared to a co-circulating R5 variant (1239, 1924, and 4102) or a heterologous R5 strain (1233). Superimposed Env structures revealed a secondary protrusion in all the phenotypically confirmed X4 V3 loops as compared to the R5 V3 loop Envs (Fig 3A - 3E). This protuberance coincided with the location of the insertion either at the tip or the base of the V3 loop (Fig. S2). The 1924 X4 V3 loop also contained a protrusion in the V3 loop in the absence of an insertion (Fig. 3E). The protuberance directly corresponded to the observed aspartic acid (D) to lysine (K) substitution at position 25 of the V3 loop compared to the R5 variant.

**Fig. 3.**
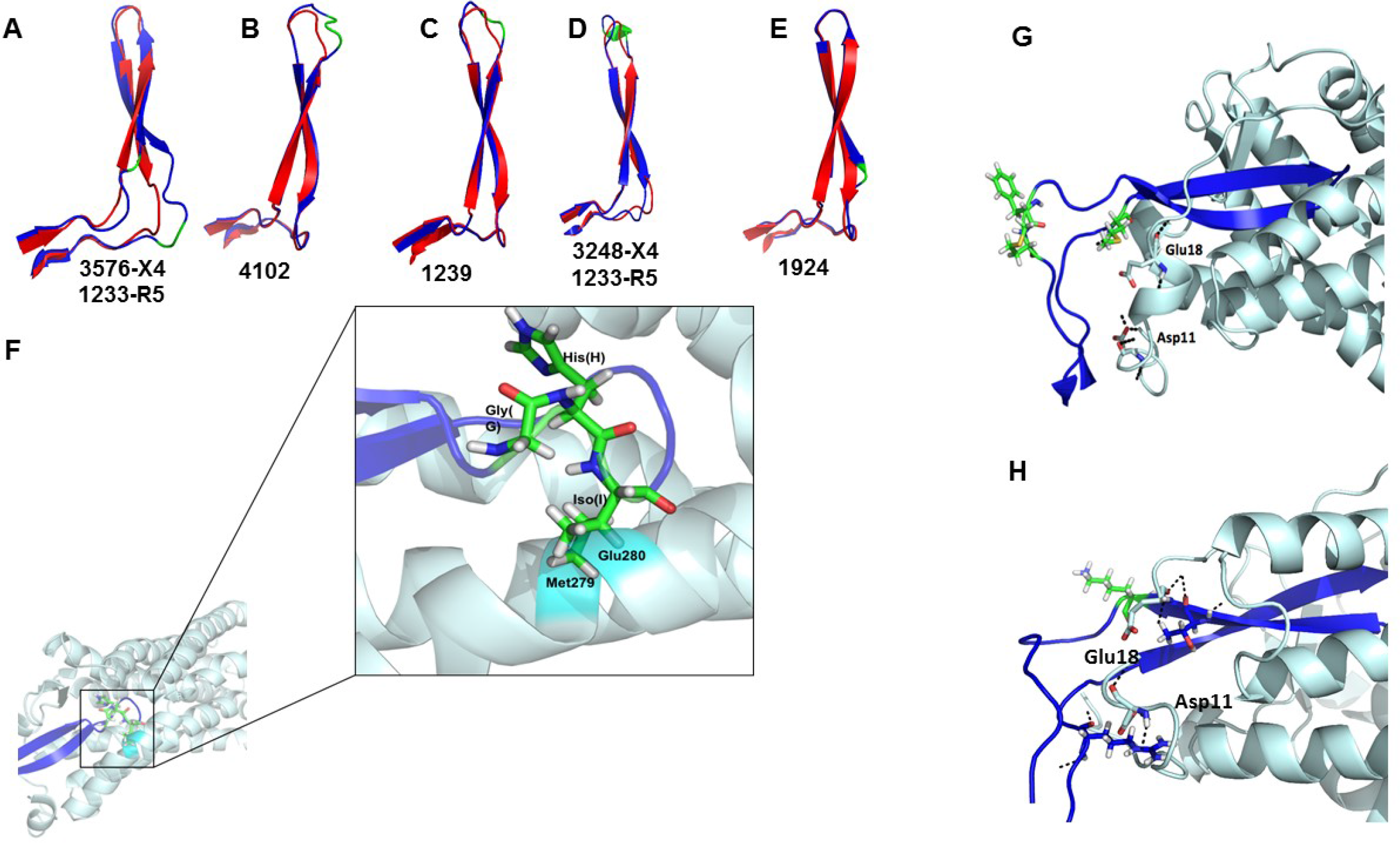
X4 V3 loops contain a protrusion that impairs CCR5 binding. Figures **A - E** depict superimposed predicted X4 (blue) and R5 (red) V3 loop structures. Location of the observed insertions and amino acid substitutions are depicted in green. ID below each predicted structure indicates the identity of the subject for the X4 V3 loop and either a co-circulating or heterologous R5 V3 loop. **(F)** Interaction of a predicted 4102-X4 V3 loop structure (purple) on the predicted 4102-R5 V3 loop – CCR5 (light blue) model. Predicted steric clash at positions 279 and 280 are highlighted in cyan. Stick configuration at the tip of the V3 loop shows the three-amino acid insertion (green) observed in the 4102 X4 Env. Interaction of 3248 **(G)** and 1924 **(H)** X4 V3 loop (purple) with the CCR5 receptor (light blue). Stick configuration shows the insertions and amino acid insertions (green) observed in these X4 Envs and amino acids Glu18 and Asp11 of the CCR5 receptor. Black dashes represent hydrogen bonds that were absent in this model compared to the 4102-R5 V3 loop and CCR5 structure (Fig. S3).

To understand the impact of these protrusions on co-receptor usage, the predicted V3 loop structure was docked with a model of the CCR5 receptor. Although an experimentally solved CCR5 structure is available, this structure is in complex with Maraviroc, an HIV entry inhibitor, which alters the Env binding pocket [37]. Thus, this solved CCR5 structure is unusable for understanding interactions with the Env V3 loop. A previously constructed structural model of CCR5 created on a CXCR4 template was used in our subsequent analysis [38]. Receptor - ligand interactions for CCR5 and a 4102 R5 V3 loop Env were predicted using Cluspro [39]. In these simulations, R5 V3 loops interacted with CCR5 in an expected manner (Fig. S3) [38].

All docking studies of CCR5 and the predicted X4 V3 loop put the X4 V3 loop in orientations that did not interact with the CCR5 receptor. In order to predict the interaction between CCR5 and an X4 V3 loop, the X4 V3 loop model was superimposed onto the predicted CCR5 - 4102 R5 V3 loop model complex (Fig. 3F - 3H). The R5-utilizing V3 loop was then removed for visual clarity. In this model, the 4102 X4 Env V3 loop crown, specifically the amino acid insertions, clashed with Methionine (M) 279 and Glutamic acid (E) 280 in the CCR5 extra-cellular loop 2 (ECL2) (Fig. 3F). On the other hand, subject 3248 X4 Env with the insertion at the base of the V3 loop eliminated two hydrogen bonds known to be important for CCR5 binding, namely V3 Arginine (R) 3 to CCR5 Aspartic acid (D) 11 and V3 R23 to CCR5 E18 (Fig. 3G) [38]. These two important hydrogen bonds were also not observed for the 1924 X4 V3 loop with the predicted protrusion (Fig. 3H). Together, these structural modeling data suggest that signature V3 sequence motifs observed in X4 strains introduce a protrusion that either sterically hinders receptor binding or eliminates important interactions with amino acids in the CCR5 N-terminal region.

Next, homology models were used to understand the impact of the X4 Envs V3 loop insertion related protrusions on V3 loop bnAb access. First, Env structures were predicted using SWISS-MODEL with Env BG505-SOSIP-gp140 as the template [40]. Next within PyMOL, the predicted models were superimposed on the crystal structure of a BG505-SOSIP in complex with either 3H+109L (PDB ID 5CEZ) or 10-1074 (PDB ID 5T3Z) [41,42]. The PGT121 precursor, 3H+109L, structure was chosen in the homology modeling because PGT121 structure has not been solved in conjunction with Env gp120 subunit. In addition, the epitope binding region for 3H+109L and PGT121 (PDB ID 4FQ1) have relatively small root-mean-square-deviation (RMSD = 1.37 Å) and similar angle of approach towards the Env (Fig. S4). Homology models revealed that the targeted V3 loop epitope was equally accessible to the PGT121 precursor and 10-1074 in the relatively resistant insertion containing X4 (4102-3_6 and 4102-3_5) and relatively sensitive insertion deficient R5 Envs (4102-61 and 4102-2_17) (Fig. 4A – 4D and Table 2). Thus, the V3 loop insertion related protrusion hinders CCR5 binding, but it does not appear to limit access to the V3 loop bnAb epitope.

**Fig. 4.**
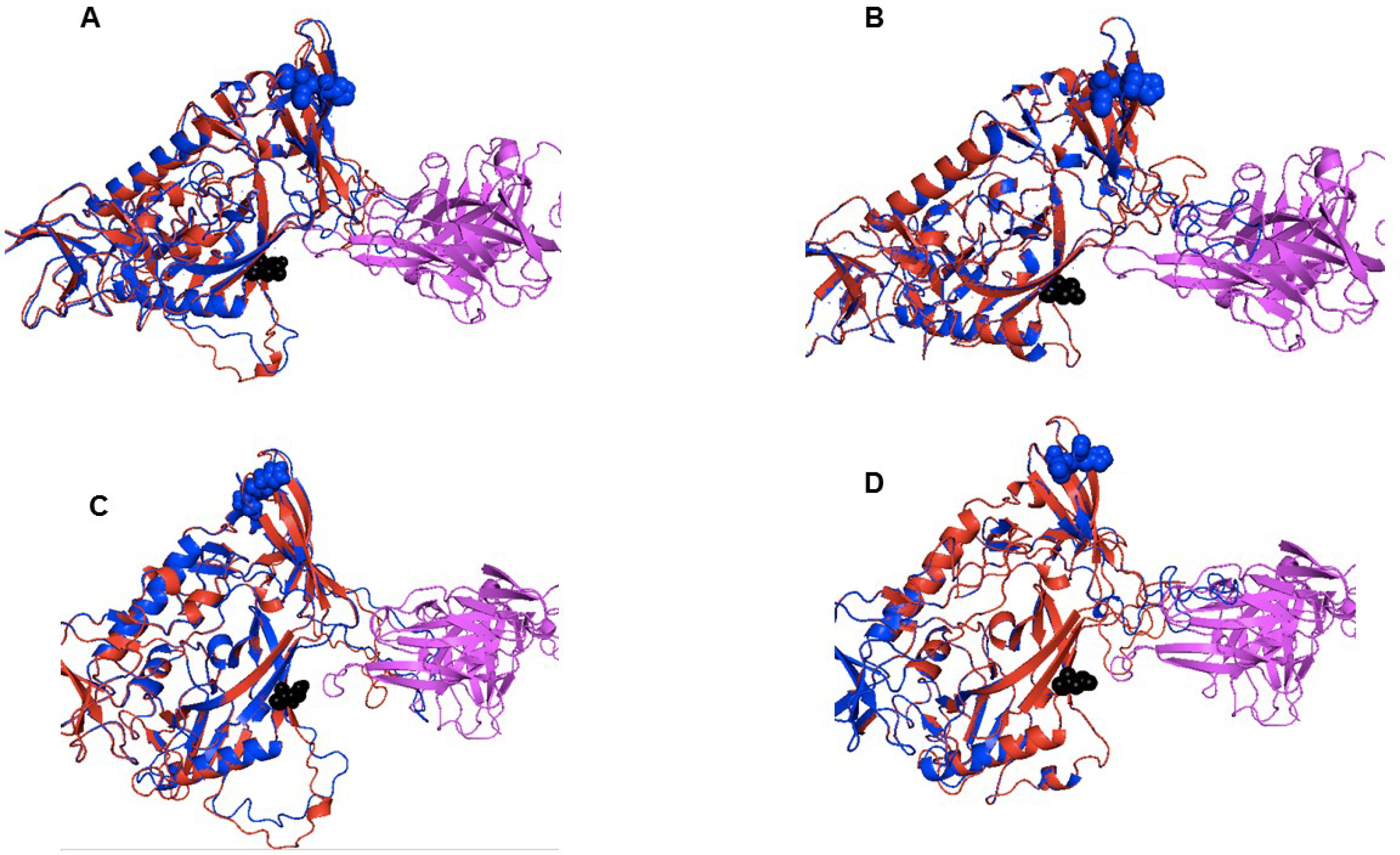
Orientation of V1 loop influences neutralization sensitivity to anti-V3 loop antibodies. Homology models of 3H+109L, a precursor to PGT121, (magenta) **(A and B)** and 10-1074 bnAb (magenta) **(C and D)** interaction with relatively sensitive R5 (4102_61 **(A and C)** and 4102-2_17 **(B and D)**) (red) and less susceptible X4 (4102-3_6 **(A and C)** and 4102-3_5 **(B and D)**) (blue) Envs. The black spheres in each structure highlight the predicted N332 site. The blue spheres show the V3 loop insertions in the X4 as compared to the R5 strains.

**Table 2.** Original and chimeric Env neutralization sensitivity and CSA

| Env ID | Phenotype | PGT121<br>(AUC) | 10-1074<br>(AUC) | 3H+109L<br>CSA (A<sup>2</sup>) | 10-1074<br>CSA (A<sup>2</sup>) |
| --- | --- | --- | --- | --- | --- |
| 4102-61 | R5 | 0.75 | 0.73 | 324.24 | 236.56 |
| 4102-3_6 | X4 | 0.24 | 0.56 | 910.51 | 869.91 |
| 4102-2_17 | R5 | 0.64 | 0.65 | 317.61 | 256.98 |
| 4102-3_5 | X4 | 0.30 | 0.33 | 1270.90 | 1150.58 |
| 3_6H-61T | R5 | 0.24 | 0.55 | 861.24 | 818.44 |
| 61H-3_6T | X4 | 0.41 | 0.64 | 769.71 | 511.05 |
| 3_5H-2_17T | R5 | 0.20 | 0.34 | 1057.91 | 771.96 |
| 2_17H-3_5T | X4 | 0.35 | 0.51 | 583.17 | 403.39 |

### Contact between the V1 loop and the bnAb impacts susceptibility

The homology models also revealed that the V1 loop of the highly sensitive Envs, pointed away from the PGT121 precursor and the 10-1074 bnAb (Fig. 4). On the other hand, the V1 loop of the relatively resistant Envs clashed with the antibodies. These structural homology models suggested that V1 loop clash impacts V3 loop bnAb sensitivity. To validate this hypothesis, we engineered chimeric Envs in which the V1-V2 loops were swapped from 4102 R5 viruses highly sensitive to PGT121 (4102-61 and 4102-2_17) and X4 variants relatively resistant to the bnAb (4102-3_6 and 4102-3_5). Chimeras contained exchanged domains from the start of the Env gene to the end of V1-V2 (labeled head (H)) and from the V1-V2 terminus to Env end (termed tail (T)). Both relatively resistant variants’ V3 loop bnAb susceptibility increased after the introduction of V1-V2 loops from the highly sensitive Envs (Table 2). In contrast, both highly sensitive R5 Envs PGT121 and 10-1074 sensitivity decreased after the introduction of the X4 V1-V2 domains. These swaps yielded Envs that were not as highly susceptible or as relatively resistant as the original non-chimeric strains. Furthermore, the V1-V2 exchanges did not switch receptor usage. In aggregate, this suggests that V1-V2 domain impact sensitivity to V3 loop bnAbs but other Env portions, such as the V3 loop, are also likely to make important contributions.

The influence of Env V1 loop orientation was further examined by estimating the contact surface area (CSA) between a predicted Env V1 structure and an antibody. In this context, higher CSA implied greater proximity of the V1 loop to the antibody and vice versa. Within PyMOL, the CSA was estimated as the sum of the solvent accessible area for the antibody and V1 loop structure individually minus the solvent accessible area for the V1 loop in complex with the antibody [43]. As expected from the predicted structures (Fig 4), the relatively resistant X4 (4102-3_6 and 4102-3_5) Envs had greater CSA as compared to the highly sensitive R5 (4102-61 and 4102-2_17) R5 Envs (Table 2). Among the original and chimeric Envs, the estimated CSA increased as neutralization PGT121 and 10-1074 AUC decreased (Fig. 5A and 5B). Thus, Envs predicted to have greater V1 loop proximity to the antibody are more neutralization resistant. To further confirm this association, 3H+109L and 10-1074 CSA was estimated for all Envs in the CATNAP database with a predicted N332 site. There was a statistically significant association between estimated CSA and bnAb sensitivity among Envs that had a detectable IC_50_ (Fig. 5C and 5D). As CSA increased, sensitivity to PGT121 and 10-1074 decreased. In aggregate, this suggests that CSA can be used to estimate V1 loop clash with an antibody, and V1 loop interference impacts neutralization susceptibility to V3 loop bnAb.

**Fig. 5.**
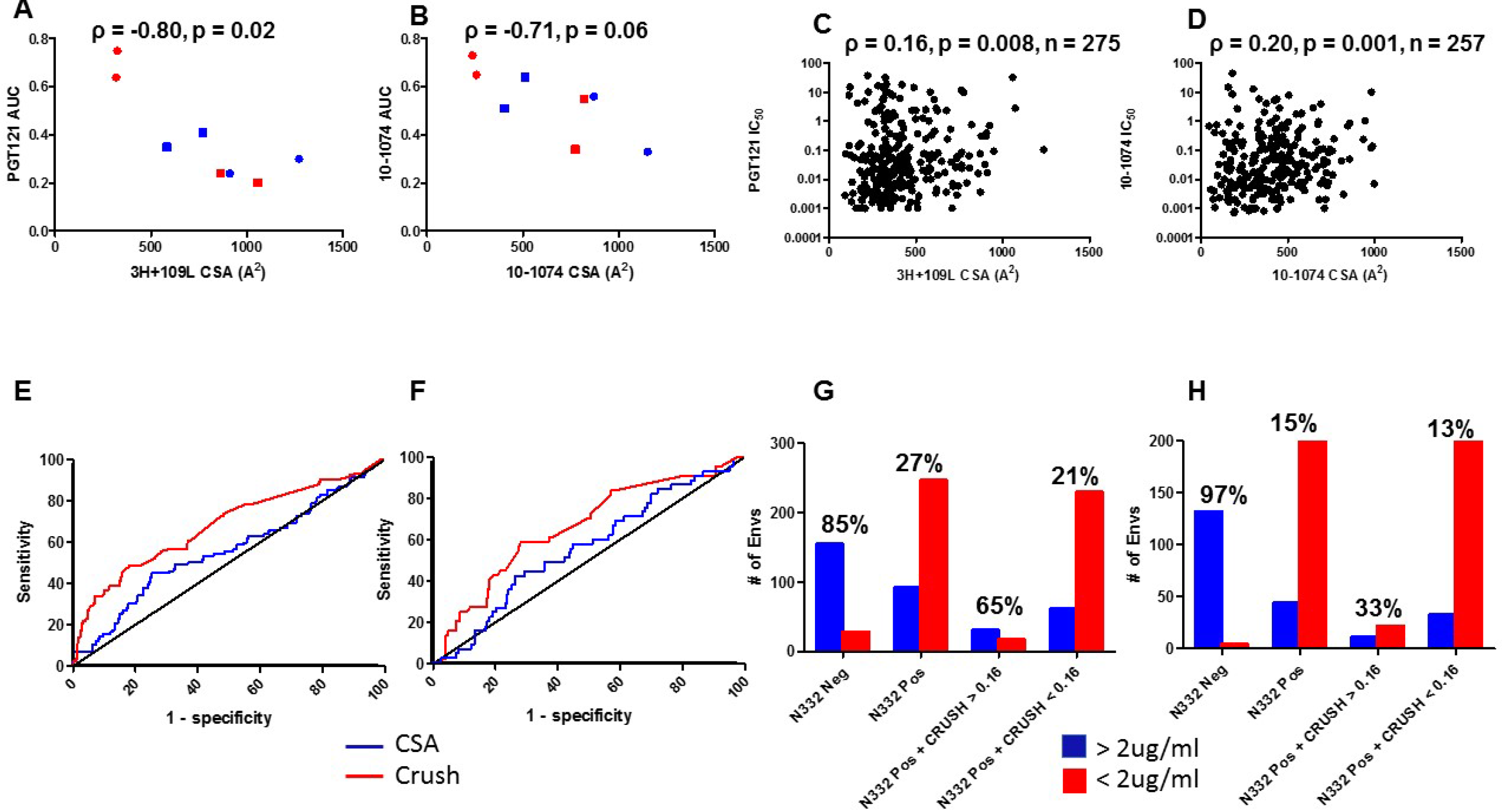
Sequence-dependent co-receptor utilization and contact between V1 loop and antibody predict neutralization sensitivity. **(A and B)** Correlation between estimated V1 loop and 3H+109L **(A)** and 10-1074 **(B)** CSA (x-axis) and PGT121 **(A)** and 10-1074 **(B)** neutralization area under the curve (AUC) (y-axis) for subject 4102 original R5 (red circles) and X4 (blue circles) Envs and chimeric R5 (red squares) and X4 (blue squares) Envs. **(C and D)** Correlation between V1 loop and 3H+109L **(C)** and 10-1074 **(D)** CSA (x-axis) and IC_50_ (y – axis) among all CATNAP Envs with detectable neutralization. **(A – D)** Graphs shows Spearman rank correlation. **(E and F)** Receiver operating curve (ROC) showing CRUSH (red) and CSA (blue) in predicting Envs with greater than versus less than 2 ug/ml PGT121 **(E)** and 10-1074 **(F)** IC_50_. The black line is the line of identity. **(G and H)** Columns depict number of Envs (y-axis) with the defined characteristic (x-axis) that had IC_50_ greater than (blue) or less than 2 ug/ml (red) for PGT121 **(G)**and 10-1074 **(H)**. Numbers above the bars denote the percent of Envs with a documented IC_50_ above 2ug/ml.

### Sequence-derived co-receptor usage but not CSA can be used to predict V3 loop bnAb sensitivity

In a future sequence-based screen, individuals harboring strains that lack a predicted glycan at the Env 332 site will likely be ineligible for V3 loop directed bnAb therapy. Numerous Envs that have the N332 site, however, still have decreased neutralization sensitivity to V3 loop bnAbs, and there are no sequence-based methods to identify these variants. We hypothesized that an algorithm that predicts co-receptor usage and estimated CSA between V1 loop and V3 loop bnAb may help distinguish less susceptible strains. The CRUSH (CoReceptor USage prediction for HIV-1) web tool was used to predict receptor utilization among all CATNAP database N332 containing Envs with neutralization data against either PGT121 or 10-1074 [20]. This highly accurate algorithm yields a probability that an input V3 Env sequence is X4. Previous clinical trials have used an IC_50_ below 2 ug/ml as an inclusion criteria for V3 loop bnAb based therapy, and thus, Envs were classified as sensitive and resistant based on this criterion [1,4]. The ability of CRUSH and V1 CSA to predict PGT121 and 10-1074 susceptibility among N332 containing Envs was initially examined by using receiver operating curves (ROC). CRUSH and V1 CSA yielded a median area under the ROC of 0.68 (95% confidence interval (CI) 0.61 – 0.75, p < 0.0001) and 0.55 (95% CI 0.48 – 0.61, p = 0.14) respectively for PGT121 (n = 338) (Fig. 5E). For 10-1074, there was also a statistically significant area under the ROC for CRUSH (median 0.66, 95% CI 0.57 – 0.75, p = 0.0006) but not for V1 CSA (median 0.55, 95% CI 0.46 – 0.65, p = 0.25) (n = 289) (Fig. 5F).

For both PGT121 and 10-1074, 175 CATNAP Envs were randomly selected as a training set to determine a CRUSH cut-off that would achieve a minimum 90% specificity for predicting an IC_50_ greater than 2 ug/ml. The remaining 163 and 114 Envs were used as a test set for PGT121 and 10-1074 respectively. In both cases, a CRUSH value more than 0.16 yielded greater than 90% specificity for the test set. This CRUSH cut-off had 93.0% (95% CI 89 – 96%) and 91% (95% CI 87 – 94%) specificity for PGT121 and 10-1074 respectively against the entire CATNAP data set. The positive predictive value (PPV) was 65% for PGT121 but only 33% for 10-1074 (Fig. 5G and 5H). This difference likely occurred because there were smaller number of CATNAP Envs with a CRUSH value greater than 0.16 with available IC_50_ data against 10-1074 (n = 33) as compared to PGT121 (n = 48), and PPV as opposed to specificity is dependent on sample composition. In both cases, however, the proportion of N332 positive Env variants with a CRUSH score greater than 0.16 had between 2 to 3 fold greater likelihood of having an V3 loop bnAb IC_50_ more than 2 ug/ml as compared to strains with predicted glycosylation at the 332 amino acid position but less than 16% probability of being X4 (Fig. 5G and 5H). Similar analysis was not conducted for CSA because it demonstrated poor test characteristics against the entire data set. In aggregate, Envs both containing the predicted primary V3 loop bnAb epitope (N332) and estimated to have greater than 16% probability of being an X4 variant had a relatively high likelihood of being relatively insensitive to PGT121 and 10-1074.

## Discussion

Passive administration of a V3 loop bnAb (10-1074) decreases plasma viremia and delays virus re-emergence in some but not all treated individuals [1,3,4]. In these trials, pre-infusion virus susceptibility impacted subsequent treatment efficacy regardless of whether 10-1074 was used as monotherapy or in combination with another bnAb. These results provide the impetus to develop techniques to screen pre-existing variants for bnAb neutralization sensitivity. In this study, we used the observation that phenotypically-confirmed CXCR4-using as compared to R5 variants are less neutralization susceptible to heterologous plasma and to V1-V2 and V3 directed bnAbs to develop such a screening test. As an application of these results, we showed that an algorithm that uses sequences to predict receptor usage identifies variants with decreased susceptibility to V3 loop bnAbs. We also developed sequence-input homology models of envelope – antibody interactions. We found that in some cases less neutralization susceptible variants have relatively large estimated contact surface between the Env V1 loop and antibody, suggesting that variable loop interference may impact bnAb potency. In aggregate, these results provide an initial sequence-based method to screen for V3 loop insensitive viruses. Although, CSA does not reliably predict neutralization sensitivity among a large set of Env variants, this sequence-dependent homology modeling provides a potential framework for developing future sequence-based tests for estimating bnAb sensitivity. This will be helpful for the various other planned bnAb clinical trials [44].

Most bnAb clinical trials have not pre-screened patients for antibody susceptibility [1–6]. Phenotypic screening using culture outgrowth techniques or Env amplification, cloning and pseudovirus production is both time and labor intensive. Importantly, phenotypic screening has low sensitivity because these methods sample a relatively small proportion of the circulating Env variants, and they may miss minor strains that are less susceptible to the bnAb under consideration [34]. It is generally agreed that a sequence-based test aimed at identifying individuals that harbor V3 loop bnAb resistant strains would first exclude those that harbor variants lacking a predicted glycan at the Env 332 site. Indeed using 2 ug/ml as a cut-off, absence of a predicted glycan at the 332 site, classified as a positive test, has around 90% and 98% specificity for identifying PGT121 and 10-1074 less sensitive strains respectively. Thus, this initial screen effectively excludes insensitive and some rare sensitive variants that lack the N332 site. Presence of the N332 glycan (a negative result), however, has a sensitivity of around 64% and 78% for PGT121 and 10-1074 respectively, suggesting that a significant proportion of the variants with the N332 site are insensitive to the V3 loop bnAbs. Our results provide novel ways to parse out these N332 containing less sensitive strains. N332 positive variants with an estimated 16% or greater probability of being an X4 strain (CRUSH score > 0.16) have around two to three fold higher likelihood of having an IC_50_ greater than 2 ug/ml as compared to the remaining N332 positive Envs. This CRUSH test, however, had relatively low PPV because only a small proportion of N332 positive strains in the CATNAP database had a CRUSH score greater than 0.16. The CATNAP database, however, may not be representative of the variants present in patients eligible for V3 directed bnAb therapy. In the CATNAP database, less than 10% of the variants were either phenotypically confirmed or predicted by sequence analysis to use the CXCR4 receptor. Natural history studies, however, estimate that often up to 50% of chronically infected individuals contain CXCR4-using viruses [45,46]. Thus, using a CRUSH cut-off as a criteria for determining the eligibility of patients for V3 loop bnAb therapy may be especially useful in settings where there is higher prevalence of X4 strains or of variants that are evolving towards CXCR4 utilization.

While specific V3 loop sequence changes are associated with R5 as compared to CXCR4-using strains, the differential neutralization susceptibility likely arose due to both V3 and non-V3 loop modifications. The V3 loop – CCR5 docking and Env – antibody homology models along with V1-V2 chimeric Env studies support this idea. We showed that a predicted protrusion near the crown of the X4 V3 loop clashed with CCR5 receptor amino acids. On the other hand, the basic amino acid substitution or protrusion at the base or middle of the X4 V3 loop eliminated important interactions with amino acids in the CCR5 N-terminal region. Prior studies have found differences in charge, hydrogen-bond donor sites, aliphatic side chain orientation, and hydrophobicity as potential explanations for co-receptor specificity [38,47–49]. This study provides a novel mechanistic understanding for the loss of CCR5 receptor usage among some exclusive CXCR4-using Envs. Env – antibody homology models predict that these V3 loop protrusions present in X4 variants, however, do not directly limit access to the epitopes important for V3 loop directed bnAb activity. Thus, the structural basis for the inability to use the CCR5 receptor does not account for decreased sensitivity to V3 loop bnAbs. Our observation that exchanging V1-V2 loops among Envs did not change co-receptor usage but it did impact sensitivity to V3 loop bnAbs further supports this notion.

The sequential co-receptor evolution from R5 to R5X4 and then X4 requires multiple sequence modifications within and outside the V3 loop [50]. We observed that Env variants with merely greater than 16% probability of being X4 had a high likelihood of having a V3 loop bnAb IC_50_ greater than 2 ug/ml. Majority of Env strains with around 16% X4 probability likely use the CCR5 and not the CXCR4 receptor to enter cells. These Env variants, however, likely contain some sequence modifications that are commonly observed among CXCR4-using strains. As opposed to only the specific V3 loop differences among R5 versus X4 strains, it is likely that the multitude of changes that occur as an Env transitions from exclusive CCR5 to only CXCR4 usage contribute to decreasing sensitivity to V3 loop bnAbs.

The V3 loop sequence changes that lead to the predicted protrusions are similar to those observed among HIV-1C, HIV-1D and simian human immunodeficiency virus (SHIV) X4 strains [14,15,51]. The HIV-1B X4 had a 2 to 3 amino acid V3 loop insertion in the same two general regions, either directly before the GPG crown or towards the base of the V3 loop. Forces promoting V3 insertions remain unclear. Neutralizing antibody (nAb) selective pressure has been associated with insertions observed in V1 thru V4 Env domains [52–54]. Strain specific V3 loop directed antibodies that bind at the crown or the base of the V3 loop are common in HIV-1 infected individuals [55,56]. BnAbs, such as PGT121 and 10-1074, also interact with residues in and around the tip of the V3 loop including the GPG crown and amino acids towards the base of the V3 loop respectively [57]. In aggregate, the similarity in the V3 loop insertions among HIV-1B, HIV-C, and HIV-1D X4 variants suggests that these highly divergent viruses are independently converging to a similar solution in response to a common selection pressure, likely nAbs. Isolating antibodies from individuals that harbor X4 strains with V3 loop insertions will provide more definitive proof for this notion.

In general, plasma samples displayed a decreased ability to neutralize Envs in the CXCR4-using as compared to the global reference panel. The global reference Env collection has been proposed as a standardized panel to evaluate neutralization capacity [29]. This panel, however, contains no CXCR4-utilizing viruses. Our results argue that CXCR4-using, especially X4 strains, should be included in a standardized Env collection for a more accurate assessment of plasma or antibody neutralization breadth and potency. This may not be important for judging the breadth and potency of potential vaccine generated antibodies because nearly all infections are initiated by R5 strains [16,45,46]. Incorporating CXCR4 variants in the standard panel to estimate neutralization capacity, however, will be important for potential future antibody-based therapeutics because chronically infected individuals often harbor CXCR4-using strains. [45,46].

Homology modeling was also used to gain a structural understanding for the linkage between differential neutralization susceptibility and co-receptor usage. The modeling and chimeric Env analysis suggested that the orientation of the V1 loop plays a role in influencing susceptibility to V3 loop directed bnAbs. Notably, these findings further confirm that Env V1-V2 loops have a major impact on sensitivity to autologous, heterologous, and now bnAbs [58,59]. Similar structure-based predictions that also incorporate interactions between different amino acids have been used to understand HIV-1 co-receptor usage [20,38]. As nAbs are introduced in the clinical arena, screening tests can use Env sequences to both predict a phenotype of interest, such as receptor usage, and develop homology structures that incorporate CSA and also electrostatic interactions between different amino acid pairs. This may yield even better sequence-based tests for predicting susceptibility to V3 loop and other bnAbs.

## Methods and Material

### Study design and samples

This study was classified as non-human subject research by the Boston University Institutional Review Board. Plasma samples were obtained from the AIDS Clinical Trials Group (ACTG) Study A5095, which was a randomized, double-blind trial assessing different ARV regimens [35]. All samples evaluated in this study were obtained before ARV therapy. One sample (SC) from a treatment-experienced individual was also available in the laboratory [31].

### Envelope isolation, virus stock production, cell lines and antibodies

Full-length Envs were amplified from each plasma sample using SGA as described previously [34]. Chimeric Envs were produced using an overlapping PCR strategy. Specific primer sequence and PCR conditions are available upon request. Amplified Envs were incorporated into a HIV-1 NL4-3 backbone to make full-length replication competent viruses using yeast gap-repair homologous recombination as described previously [60]. Briefly, virus stocks were generated by co-transfecting human epithelial kidney (HEK) 293T with a plasmid containing the Env of interest and a helper plasmid. Virus stocks were passaged in peripheral blood mononuclear cells (PBMCs) obtained from HIV-1 seronegative donors for a maximum of 7 days. Viral titers were determined using TZM-bl cells as described previously [34]. All cell lines and antibodies were obtained from the NIH AIDS Reference Reagent Program.

### Genotype prediction, co- receptor usage and sequence analysis

Each SGA Env co-receptor phenotype was predicted from the V3 loop sequence using either WebPSSM [21] or Geno2Pheno at a false predication rate of 5% [22]. Phenotypic co-receptor usage was determined by infecting TZM-bl cells in the presence or absence of TAK779 and/or AMD3100 as described previously [34]. All assays were performed along with Envs with known co-receptor phenotype. No Env showed replication in the presence of both inhibitors, and thus this confirmed that the viruses only entered cells by using one or both of the receptors. Env amplified products were cleaned using ExoSap IT (Affymetrix), and sequences were determined using Sanger sequencing.

### Neutralization assay

Neutralization sensitivity was tested by assessing infection of TZM-bl cells in the presence or absence of serial dilution of plasma or bnAb as described previously [34]. All plasma was heat inactivated at 56 for 1hr to prevent subsequent complement mediated inhibition. Area under the curve was calculated as described [36]. None of the plasma or antibody demonstrated neutralization against HIV-1 Env deleted pseudovirions with vesicular stomatitis virus G envelope protein, suggesting there was no non-specific inhibition.

Neutralization against the global reference Env and the CXCR4-using Env panel was assessed only at one plasma dilution (1:50). BP score were calculated using the equation as described previously [30].

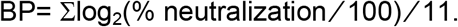

A score of 0 represents no neutralization and a score of 1 represents 100% neutralization. Heat maps were generated using the Los Alamos HIV sequence database heat map tool (https://www.hiv.lanl.gov/). All heat maps used hierarchical clustering with the Euclidean distance method.

### Structural modeling and docking

Models of X4- and R5-utilizing V3 loops were produced using Rosetta software made available by Robetta Structural Prediction Server online [61]. Model 1, the best model based on ProQ2 rank, was selected for each V3 loop. Docking of the CCR5 chemokine receptor with R5 and X4 V3 loops were done using Cluspro [39]. All superimpositions were done using PyMOL software (Schrödinger LLC version 2.2.2). Table S4 lists the HIV-1 template chosen by the server to predict the V3 loop structure.

Env homology models were generated using SWISS MODEL with BG505 SOSIP.664 as the user input template [40]. Env homology models were superimposed with the PGT121 precursor, 3H+109L (PDB ID 5CEZ) or 10-1074 (PDB ID 5T3X) using PyMOL software (Schrodinger LLC version 2.2.2). Contact surface area was generated using an open source code available at https://pymolwiki.org/index.php/Contact_Surface.

### Statistical Analysis

Comparisons were done among all Los Alamos CATNAP database Envs with previous neutralization data against specific bnAbs. Envs with phenotypically-defined CXCR4-usage were compared to those with exclusive CCR5 usage. Detectable versus undetectable neutralization sensitivity was defined based on the presence of an estimated IC_50_ less than or greater than the highest tested antibody concentration respectively. Envs with undetectable IC_50_ were assigned a value of 100 ug/ml for statistical comparisons. Env receptor usage and CSA was estimated for all Envs with available sequence data, a predicted glycan at the Env 332 site, and neutralization data against 10-1074 and PGT121. Co-receptor usage was predicted using CRUSH (CoReceptor USage prediction for HIV-1) web tool [20].

Comparisons between groups containing independent data points or matched samples were done using the Mann-Whitney test and the Wilcoxon matched-pairs test respectively. Frequency differences were examined using two-sample test of proportions. Associations were estimated using Spearman rank correlations. ROC were estimated by separating Envs into groups with IC_50_ greater and less than 2 ug/ml. Statistical analyses were done using GraphPad Prism 5 (version 5). All p-values are based on two sided tests.

CRUSH and CSA prediction characteristics were also examined using logistic regression. In this analysis, CSA and CRUSH were examined as predictors of having an IC_50_ greater than or less than 2 ug/ml. Results were assessed using 2-fold cross validation and repeated 1000 times. The results using this analysis were not significantly different as compared to the ROC evaluation. In addition, multi-variate logistic regression analysis with both CSA and CRUSH did not improve prediction.

## Acknowledgements

We would like to thank all of the ACTG Study A5095 participants. We thank Laura White and Wenqing Jiang (Providence/Boston Center for AIDS Research biostatistics core) for helpful discussions. We thank Budhi Sagar for technical insights.

## Supplementary Materials

Fig. S1. Correlation between IC_50_ and neutralization area under the curve.

Fig. S2. Predicted amino acid Env sequence alignment.

Fig. S3. Predicted interaction between CCR5 and a R5 V3 loop.

Fig. S4. Predicted PGT121 and 3H+109L Env complex.

Supplementary Table 1. Samples with R5 only and dual-mixed virus population.

Supplementary Table 2. Envelopes in the CXCR4-using Env panel.

Supplementary Table 3. Variants in the primary X4 and R5 Env panel.

Supplementary Table 4. Templates for homology modeling for X4 and R5 V3 loops.

## Notes

The authors have declared that no conflict of interest exists

